# Opposing motors provide mechanical and functional robustness in the human spindle

**DOI:** 10.1101/2021.03.02.433652

**Authors:** Lila Neahring, Nathan H. Cho, Sophie Dumont

## Abstract

At each cell division, the spindle self-organizes from microtubules and motors. How the spindle’s diverse motors, often acting redundantly or in opposition, collectively give rise to its emergent architecture, mechanics, and function is unknown. In human spindles, the motors dynein and Eg5 generate contractile and extensile stress, respectively. Inhibiting dynein or its targeting factor NuMA leads to unfocused, turbulent spindles and inhibiting Eg5 leads to monopoles, yet bipolar spindles form when both are inhibited together. What, then, are the roles of these opposing motors? Here we generate NuMA/dynein- and Eg5-doubly inhibited spindles that not only attain a typical metaphase shape and size, but also undergo anaphase. However, these spindles have reduced microtubule dynamics and are mechanically fragile, fracturing under force. Further, they exhibit lagging chromosomes and dramatic left-handed twist at anaphase. Thus, while these opposing motor activities are not required for the spindle’s shape, they are essential to its mechanical and functional robustness. Together, this work suggests a design principle whereby opposing active stresses provide robustness to force-generating cellular structures.

## Introduction

At each cell division, the spindle self-organizes from dynamic microtubules, crosslinkers, and motors (Elting et al., 2018; McIntosh et al., 2012). Together, these molecular-scale force generators give rise to a cellular-scale structure with emergent properties such as a steady-state shape in metaphase and the ability to accurately segregate chromosomes at anaphase. The mammalian spindle’s molecular components have been extensively cataloged (Neumann et al., 2010), and the biophysical properties of many individual motors are now known. However, it remains poorly understood how combinations of motor activities—many of which act redundantly or in opposition to each other—give rise to the mammalian spindle’s emergent architecture, mechanics, and function.

The motors Eg5 and dynein are key determinants of spindle architecture. Both generate directional forces between pairs of microtubules that they crosslink, building distinct cellular-scale motifs that coexist in the spindle’s microtubule network. Eg5 is a bipolar homotetrameric kinesin-5 that slides antiparallel microtubules apart, generating extensile stress in the spindle and maintaining pole separation (Blangy et al., 1995; Kapitein et al., 2005; Roostalu et al., 2018; Shimamoto et al., 2015). Conversely, dynein is recruited to microtubule minus ends by its targeting factor NuMA, where it generates contractile stress by carrying minus end cargoes towards the minus ends of neighboring microtubules (Figure 1A) (Foster et al., 2015; Gaglio et al., 1996; Hueschen et al., 2017). These motors have opposing loss-of-function phenotypes that are deleterious for the dividing cell. When Eg5 is inhibited, spindles form as monopoles with minus ends clustered into a single aster (Mayer et al., 1999), whereas NuMA or dynein deletion leads to turbulent, disordered spindles with no steady-state shape and with microtubule bundles extending against the cell cortex (Hueschen et al., 2019).

**Figure 1.**
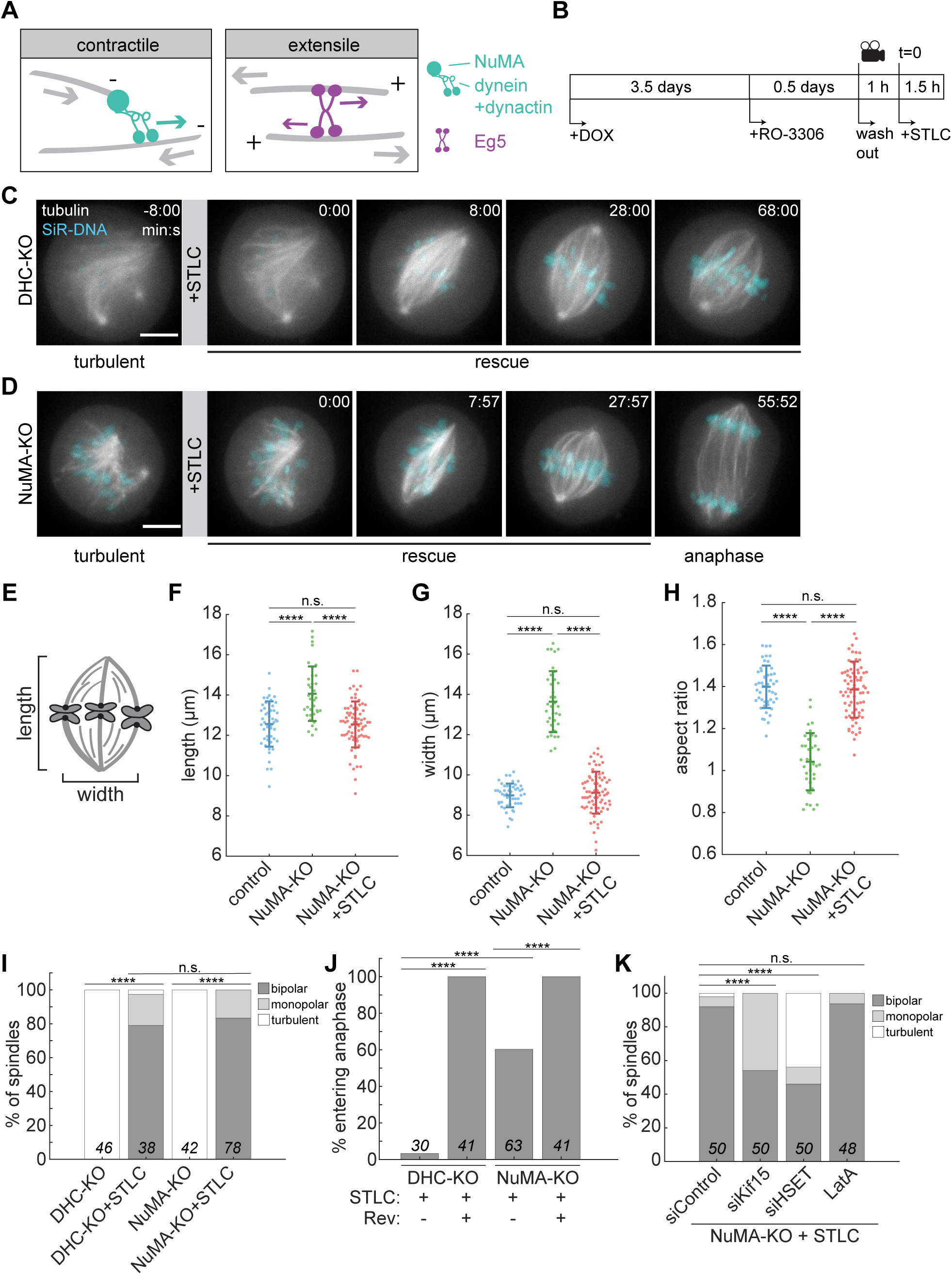
Eg5 Inhibition Allows Turbulent Spindles to Recover Bipolarity and Progress to Anaphase. (A) Schematic illustrations of contractile microtubule (gray filament) minus end clustering by dynein, dynactin, and NuMA (left, green), and extensile sliding of antiparallel microtubules by Eg5 (right, purple) in the human spindle. Dynein/dynactin, targeted to minus end cargoes by NuMA, walks towards microtubule minus ends (denoted by “-“). Eg5 walks towards microtubule plus ends (denoted by “+”). Direction of motor stepping is indicated by green and purple arrows, and contractile and extensile stresses are indicated by gray arrows. (B) Schematic diagram of opposing motor (NuMA/dynein and Eg5) inhibition experiment in human spindles. Cas9 expression was induced by doxycycline addition (+DOX) for 4 days to knock out dynein heavy chain or NuMA. Cells were synchronized in G2 (with Cdk1 inhibitor RO-3306) for 0.5 days before imaging, released into mitosis, and Eg5 was acutely inhibited during imaging with 5 µM STLC. See also Figure S1. (C) Representative timelapse confocal images of an RPE1 DHC-KO cell stably expressing GFP-tubulin (gray, maximum intensity projection of 5 planes) with SiR-DNA labeling chromosomes (cyan, single plane), starting as a turbulent spindle. After 5 µM STLC addition to inhibit Eg5 (time 0:00), the turbulent spindle recovers bipolarity, but does not progress to anaphase. Scale bar = 5 µm. (D) Representative timelapse confocal images of an RPE1 NuMA-KO cell stably expressing GFP-tubulin (gray, maximum intensity projection of 5 planes) and mCherry-H2B (cyan, single plane), starting as a turbulent spindle. After 5 µM STLC addition to inhibit Eg5 (time 0:00), the turbulent spindle recovers bipolarity and progresses to anaphase. Scale bar = 5 µm. (E) Schematic illustration of spindle length and width measurements. (F) – (H) Length (F), width (G), and aspect ratio (length/width; (H)) of control (-DOX), turbulent NuMA-KO, and bipolar NuMA-KO+STLC spindles. Spindle dimensions were measured after establishment of bipolarity (control, NuMA-KO+STLC) or 45 min after the start of imaging (NuMA-KO). Data in (F)-(H) include the same 49 (control), 36 (NuMA-KO), and 75 (NuMA-KO+STLC) spindles pooled from ≥ 3 independent experiments. ****, p < 0.00005; n.s. = not significant, two-sample t-test. Lines represent mean ± s.d. (I) Outcomes 90 min post-STLC addition to NuMA- and DHC-KO turbulent spindles. Without STLC addition, DHC-KO and NuMA-KO spindles remain turbulent. After STLC addition, most spindles establish bipolarity. (J) Percentage of bipolar spindles entering anaphase within 90 min of STLC addition, with and without 500 nM of the MPS1 inhibitor reversine to bypass the SAC. DHC-KO+STLC cells enter anaphase after reversine addition, consistent with DHC-KO+STLC cells experiencing a SAC-dependent metaphase arrest. (K) Spindle outcomes in NuMA-KO cells 90 min after STLC addition, with luciferase (Control), Kif15, or HSET RNAi knockdown or 500 nM latrunculin A to disrupt actin. Kif15 and HSET are required for turbulent spindles to recover bipolarity in the absence of NuMA and Eg5, and F-actin is not. See also Figures S1E and S1F. For (I)-(K), number of spindles is indicated on each bar; cells pooled from ≥3 independent experiments. ****, p < 0.00005, n.s. = not significant, Fisher’s exact test.

Despite their importance to spindle architecture, when dynein and Eg5 are co-depleted, human spindles form as typical bipoles (Florian and Mayer, 2012; Tanenbaum et al., 2008; van Heesbeen et al., 2014). Similar phenomena have been reported in yeast, *Drosophila*, *Xenopus laevis* extract, and pig spindles when the homologous kinesin-5 and the dominant end-clustering motor (dynein or a kinesin-14) are inhibited (Ferenz et al., 2009; Mitchison et al., 2005; Rincon et al., 2017; Saunders and Hoyt, 1992; Sharp et al., 1999). These observations suggest that the balance of contractile and extensile stress in the spindle is more important than the specific magnitude of these stresses. This raises the question: what are the functions of opposing, energy-consuming motor activities in the spindle if the same structure can be formed without them? Previous work in *Xenopus* extract spindles has suggested a role for opposing activities of dynein and Eg5 in establishing the spindle’s microtubule organization, mechanical integrity, and heterogeneity (Brugues et al., 2012; Mitchison et al., 2005; Takagi et al., 2019), but it is unknown if this applies to other spindles, whose architectures differ and whose mechanics are challenging to probe. Furthermore, dynein performs multiple functions at the kinetochore in addition to its role in minus end clustering (Howell et al., 2001; Raaijmakers and Medema, 2014), complicating the interpretation of dynein- and Eg5-doubly inhibited phenotypes in metaphase and limiting their study in anaphase. Thus, while antagonistic contractile and extensile stress generation is a highly conserved feature of the spindle, its mechanical and functional roles throughout the spindle’s lifetime remain unclear.

Here, we show that while the opposing motor activities of NuMA/dynein and Eg5 are not required to build the human spindle, they are instead essential to its robustness—the spindle’s ability to tolerate mechanical and biochemical fluctuations while maintaining its integrity and functional accuracy. Without these opposing motor activities, we find that spindles are more fragile when mechanically challenged in metaphase, and highly twisted and error-prone in anaphase. More broadly, these findings suggest a design principle by which opposing active force generators make self-organizing cellular structures robust.

## Results

### Eg5 Inhibition Allows Turbulent Spindles to Recover Bipolarity and Progress to Anaphase

To generate human spindles lacking the opposing motor activities of NuMA/dynein and Eg5, we used an inducible CRISPR knockout (KO) approach (McKinley and Cheeseman, 2017) to delete either dynein heavy chain (DHC) or NuMA in RPE1 cells (Figures S1A-S1C). This results in chaotic, turbulent spindles (Hueschen et al., 2019), in contrast to the steady-state barrel-shaped spindles resulting from RNAi depletion of DHC (Tanenbaum et al., 2008). We induced Cas9 expression for 4 days to knock out DHC or NuMA, synchronized cells with the Cdk1 inhibitor RO-3306, and released cells into mitosis before live imaging labeled microtubules and chromosomes to ensure that turbulent spindles did not accumulate defects during an extended mitotic arrest (Figure 1B). As previously reported, spindles with DHC and NuMA deleted exhibited a very similar turbulent phenotype, consistent with NuMA and dynein’s acting as a complex to cluster microtubule minus ends (Hueschen et al., 2019; Hueschen et al., 2017). After confirming knockout in each cell via the turbulent phenotype, we acutely inhibited Eg5 with S-trityl-L-cysteine (STLC), leading both DHC-KO and NuMA-KO spindles to recover into steady-state metaphase bipoles (Figures 1C and 1D; Movie S1). Many doubly inhibited spindles exhibited local defects that dynamically arose and repaired (Figure S1D), but in contrast to expanded turbulent NuMA-KO spindles, global metaphase spindle shape and size was indistinguishable from controls (Figures 1E-1H). The rescue of bipolarity was highly reproducible and dependent on Eg5 inhibition (Figure 1I). A key advantage of this experimental system was that many (60.3%) of the NuMA-KO+STLC spindles progressed to anaphase within 90 min of STLC addition. In contrast, few (3.3%) of the DHC-KO+STLC spindles did (Figure 1J). Consistent with dynein’s NuMA-independent role at the kinetochore in silencing the spindle assembly checkpoint (SAC) (Gassmann et al., 2010; Howell et al., 2001), bypassing the SAC using the MPS1 inhibitor reversine caused almost all NuMA- and DHC-KO+STLC cells to enter anaphase (Figure 1J). Thus, we used NuMA-KO cells for the remainder of our experiments to isolate dynein’s minus end clustering role from its kinetochore functions, enabling us to investigate the roles of opposing motors in both metaphase and anaphase.

To probe the mechanisms that bipolarize turbulent spindles in the absence of NuMA/dynein and Eg5, we depleted additional candidate motors in the context of our live-imaged double inhibition experiment. We found that the kinesins Kif15 and HSET are both required to establish bipolarity (Figure 1K; Movie S2). Partially depleting Kif15 in doubly inhibited spindles led to the formation of more monopoles (46%; Figures 1K, S1E, and S1F), consistent with its role in generating extensile stress (Sturgill and Ohi, 2013; Tanenbaum et al., 2009; van Heesbeen et al., 2014; Vanneste et al., 2009). Conversely, depleting HSET caused more spindles to remain turbulent (44%), indicating that it performs contractile minus end clustering. This is consistent with HSET’s roles in aster formation *in vitro* (Mountain et al., 1999; Norris et al., 2018) and centrosome clustering in cancer cells (Kwon et al., 2008). F-actin was not required to focus minus ends during bipolarization (Figure 1K), despite its importance in centrosome clustering (Kwon et al., 2008). In sum, upon Eg5 inhibition, the partially redundant contractile and extensile motors HSET and Kif15 allow turbulent NuMA-KO spindles to establish bipolarity, attain a typical metaphase size, and undergo anaphase. Thus, these doubly inhibited spindles provide a system for probing the contributions of opposing motors to spindle mechanics and function independently of spindle shape and size.

### Microtubule Organization and Dynamics are Disrupted in Doubly Inhibited Spindles

Although NuMA/dynein and Eg5 are not required for spindle shape, both of these motors contribute to the continuous transport of non-kinetochore microtubules in the human spindle (Lecland and Luders, 2014). Thus, to determine the role of these opposing motor activities, we first tested the hypothesis that they are required for the spindle’s internal organization and dynamics. We examined microtubule organization by quantifying the distribution of tubulin intensity along the spindle’s pole-to-pole axis. As expected (Crowder et al., 2015), control cells had strongest tubulin intensity near the two poles and lower intensity near the spindle equator. In contrast, tubulin intensity was more uniform in NuMA- and Eg5-doubly inhibited spindles (Figures 2A and 2B). This pattern was not due to a difference in chromosome alignment, as the intensity profile of the DNA stain Hoechst overlapped between the two conditions (Figure 2C). Thus, doubly inhibited spindles have altered microtubule organization, indicating that microtubule transport, nucleation, and/or length regulation in the spindle is disrupted without NuMA and Eg5.

**Figure 2.**
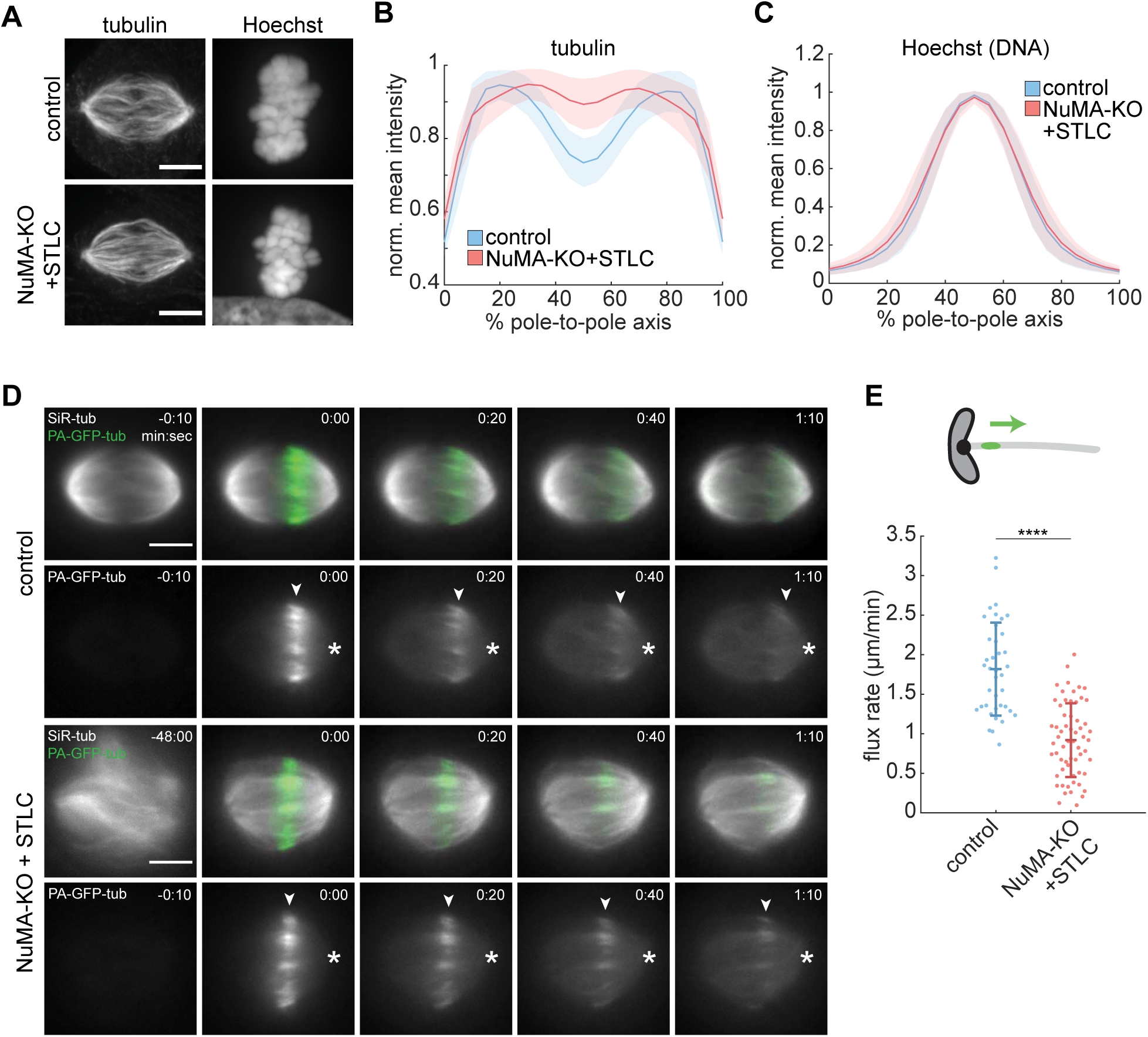
Microtubule Organization and Dynamics Are Disrupted in Doubly Inhibited Spindles. (A) Representative immunofluorescence images (maximum intensity projections) of control and NuMA-KO+STLC RPE1 cells, stained for tubulin (left) and with Hoechst (right). Scale bar = 5 µm. (B-C) Distributions of mean tubulin (B) and Hoechst (C) intensity at each point along the spindle’s pole-to-pole axis, quantified from sum intensity projections of immunofluorescence images and normalized to the maximum value in each spindle (see Methods). Doubly inhibited spindles have defects in microtubule organization. (B) and (C) include the same 335 control and 336 NuMA-KO+STLC cells pooled from 8 independent experiments. Plots represent mean ± s.d. (D) Representative timelapse widefield images of RPE1 control and NuMA-KO+STLC cells stably expressing photoactivatable (PA)-GFP-tubulin (green), co-labeled with 100 nM SiR-tubulin (gray) and photomarked near the spindle equator (t = 0:00). The PA-GFP-tubulin channel alone is shown below the merged images. Arrowheads track the photomark position, and asterisks mark the spindle pole. Scale bars = 5 µm. (E) Poleward flux rates in control and NuMA-KO+STLC cells, showing reduced microtubule transport in doubly inhibited spindles. Each dot represents an individual k-fiber. *n* = 39 k-fibers pooled from 14 cells in 1 experiment (control), *n* = 61 k-fibers pooled from 25 cells in 5 independent experiments (NuMA-KO+STLC). ****, p < 0.00005, two-sample t-test. Lines represent mean ± s.d.

We next asked whether the altered spatial distribution of microtubules in doubly inhibited spindles was associated with perturbed microtubule dynamics. We expressed photoactivatable-GFP-tubulin in NuMA-KO cells, co-labeled spindles with SiR-tubulin, and photoactivated stripes near the metaphase plate (Figure 2D; Movie S3). Tracking photomark movements on individual kinetochore-fibers (k-fibers) revealed that the k-fiber poleward flux rate was halved in doubly inhibited spindles compared to controls (0.9 ± 0.5 µm/min compared to 1.8 ± 0.6 µm/min; Figure 2E). While outward sliding by Eg5 drives microtubule flux in *Xenopus laevis* extract spindles (Miyamoto et al., 2004), k-fiber flux in mammalian spindles is thought to be largely powered by mechanisms other than Eg5 (Cameron et al., 2006; Ganem et al., 2005; Steblyanko et al., 2020). However, our findings indicate that NuMA and Eg5 are together key to microtubule flux in the human spindle. Thus, while other motors can establish the spindle’s global shape and size without the opposing forces generated by NuMA/dynein and Eg5 (Figure 1), they cannot recapitulate its locally specialized microtubule organization and dynamics.

### Doubly Inhibited Spindles are Structurally Unstable in Response to Mechanical Force

We next tested the hypothesis that opposing motors contribute to the spindle’s ability to maintain its structure under force. Loss of opposing NuMA/dynein and Eg5 activities could give rise to mechanical defects through reduced microtubule organization and dynamics (Figure 2), or through changes to the spindle’s material properties as a result of altered local force generation. To probe the mechanics of NuMA- and Eg5-doubly inhibited spindles, we reproducibly confined metaphase cells in PDMS devices (Le Berre et al., 2012), forcing them into a flattened 5 μm-high geometry (Figure 3A). Doubly inhibited spindles exhibited a different characteristic response to confinement than controls, both in their deformation over time and in their loss of structural integrity (Figure 3B; Movie S4). Although all spindles widened and lengthened during confinement, controls reached a new steady-state size after the first few minutes (Dumont and Mitchison, 2009) while doubly inhibited spindles continued to expand, failing to reach a new steady-state size in our observation period. During initial expansion, spindles in both conditions widened similarly but doubly inhibited spindles lengthened more slowly, consistent with a role of NuMA/dynein in spindle elongation (Guild et al., 2017). However, doubly inhibited spindles continued to grow in both dimensions throughout the perturbation, ultimately surpassing the new steady-state mean length and width of controls (Figures 3C and 3D). As another metric of spindle shape evolution, we calculated the 2-D correlation coefficient between binarized masks of the same spindle at multiple timepoint pairs. The shape correlation of doubly inhibited spindles was lower than that of controls at increasing lag times, and exponential fits revealed that shape correlation decayed to a lower minimum value for doubly inhibited spindles (Figure 3E). Thus, under force, doubly inhibited spindles not only deform more but have a weaker “shape memory” than controls.

**Figure 3.**
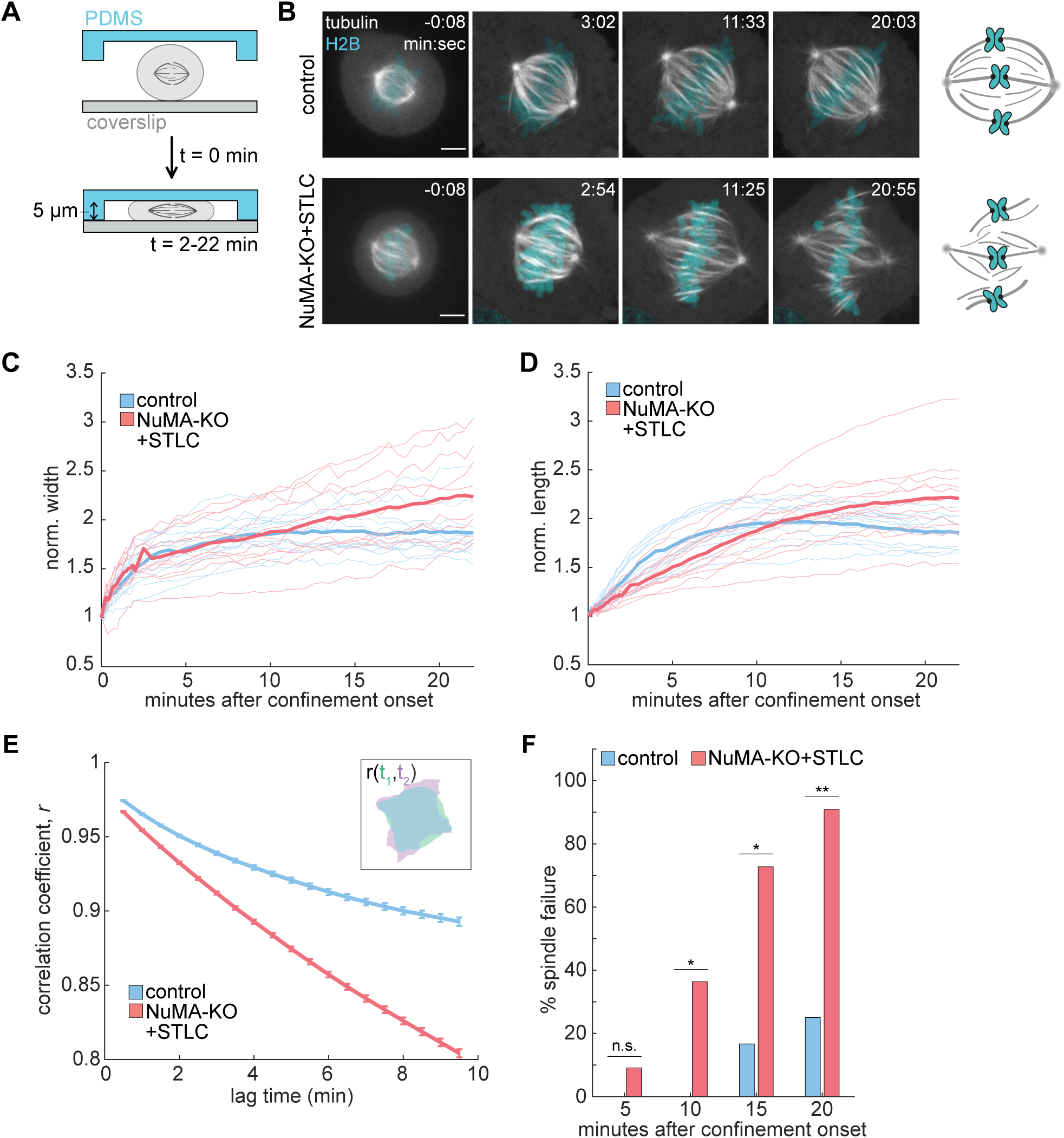
Doubly Inhibited Spindles are Structurally Unstable in Response to Mechanical Force. (A) Schematic illustration of cell confinement experiment to probe spindle mechanical robustness. Confinement to 5 μm was applied over a period of 2 min, and the confined geometry was sustained for an additional 20 min. (B) Timelapse confocal images of control and NuMA-KO+STLC RPE1 cells stably expressing GFP-tubulin (gray) and H2B (cyan) during confinement (begins at t = 0:00). K-fibers detach from poles in the doubly inhibited spindle, while the control spindle remains intact, as cartooned (right). Scale bars = 5 µm. (C-D) Spindle width (C) and length (D) during confinement of control and NuMA-KO+STLC RPE1 cells, normalized to the initial length and width of each spindle. Mean values shown in bold lines. *n* = 12 control and 11 NuMA-KO+STLC cells, pooled from 5 and 4 independent experiments, respectively. (E) Mean ± s.e.m. of spindle shape correlation coefficient between all pairs of two binary, segmented frames (green t_1_, purple t_2_ in inset), as a function of the time elapsed between the two frames (t_2_-t_1_). Shape correlation was fit to the exponential function *r* = *a* * 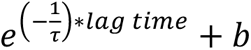, where *b* = 0.87 for controls and *b* = 0.58 for NuMA-KO+STLC. Analysis includes the same cells as (C-D). (F) Percentage of spindles that structurally fail under confinement, defined qualitatively as a loss of continuity between k-fibers and poles. Doubly inhibited spindles begin to fail earlier, and fail more frequently, than controls. *, p < 0.05; **, p < 0.005; n.s., not significant; Fisher’s exact test. Analysis includes the same cells as (C-E).

Strikingly, the impaired ability of doubly inhibited spindles to stabilize their shapes was associated with increased structural failure. By 20 minutes after confinement onset, k-fibers had detached from poles in 91% of doubly inhibited spindles, compared to 25% of controls (Figures 3B and 3F). Although control spindle poles can split during sustained confinement (Lancaster et al., 2013), failure in doubly inhibited spindles began sooner and occurred more frequently (Figure 3F). Moreover, the mode of failure qualitatively differed between the two conditions: while detached k-fibers in control spindles remained clustered into acentrosomal foci, k-fibers in doubly inhibited spindles splayed as individual bundles. Thus, while unperturbed doubly inhibited spindles maintain a similar geometry to controls (Figures 1F-1H), their reduced structural integrity becomes evident upon mechanical challenge. Together, the larger deformation, lack of new steady-state establishment, and structural fragility of doubly inhibited spindles under force indicate that NuMA and Eg5 are essential to the spindle’s mechanical robustness.

### Spindles with Reduced Opposing Motor Activity Exhibit Twist and Functional Defects in Anaphase

Given that NuMA/dynein and Eg5 are together required for the metaphase spindle’s internal organization, dynamics (Figure 2) and mechanical robustness (Figure 3), we next sought to determine whether they are important to anaphase spindle structure and function. Our finding that many NuMA-KO+STLC spindles undergo anaphase (Figure 1) allowed us to address this question. In the first 3 minutes of anaphase doubly inhibited spindles elongated, and chromosomes segregated, at rates indistinguishable from controls (Figures 4A-4C, S2A and S2B; Movie S5). However, in doubly inhibited cells spindle elongation and chromosome segregation continued at these rates for extended durations, causing spindle poles to often hit the cortex and chromosomes to segregate to greater distances (Figures 4C, S2C and S2D). Although cortical NuMA/dynein complexes generate anaphase pulling forces in other systems (Aist et al., 1993; Grill et al., 2001), and although Eg5 has been reported to contribute to outward sliding during human spindle elongation (Vukusic et al., 2019), our results indicate that NuMA- and Eg5-doubly inhibited spindles are not deficient in elongation but instead over-elongate in anaphase. Thus, either doubly inhibited spindles are subject to increased outward forces in anaphase, or they resist them less strongly.

**Figure 4.**
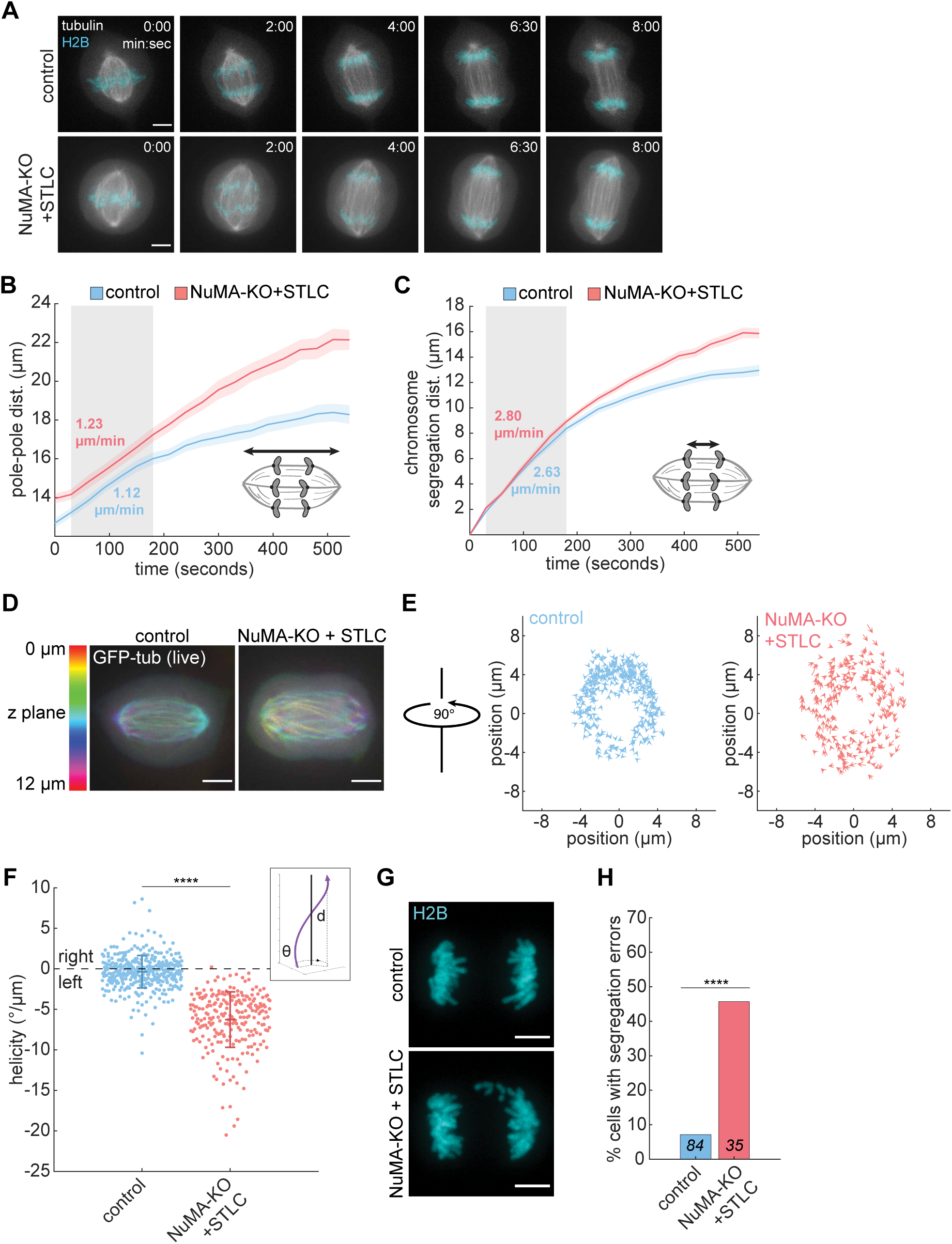
Spindles with Reduced Opposing Motor Activity Exhibit Twist and Functional Defects in Anaphase. (A) Representative timelapse confocal images of control and NuMA-KO+STLC RPE1 cells, stably expressing GFP-tubulin (gray) and H2B (cyan), during anaphase (begins at t = 0:00). Images represent a single z-plane. Scale bars = 5 µm. (B) Spindle pole-to-pole distance during anaphase (aligned to anaphase onset at t = 0), quantified from timelapse imaging of control and NuMA-KO+STLC RPE1 cells stably expressing GFP-tubulin and mCherry-H2B. Spindles initially elongate at indistinguishable rates (mean rates calculated over gray boxed area), but ultimately elongate more in doubly inhibited spindles. Lines and shaded regions indicate mean ± s.e.m. of 20 cells (control) or 18 cells (NuMA-KO+STLC) pooled from 4 independent days. See also Figure S2. (C) Distance between the two segregating chromosome masses in anaphase (anaphase onset at t = 0), same cells as (B). Chromosomes initially segregate at indistinguishable rates (mean rates calculated over gray boxed area), but segregate a greater total distance in doubly inhibited spindles. Lines and shaded regions indicate mean ± s.e.m. See also Figure S2. (D) Representative confocal images of GFP-tubulin-labeled control (left) and NuMA-KO+STLC (right) RPE1 anaphase cells, showing a single timepoint from live imaging. Spindles are colored by z-plane. Scale bars = 5 µm. (E) Spindle pole end-on views (90° rotation compared to view in (D)) of tracked interpolar microtubule bundles in control and NuMA-KO+STLC anaphase spindles. Arrow vectors (length and direction) represent the displacement of each bundle per µm traversed along the pole-to-pole axis, moving towards the viewer. *n* = 370 bundles, pooled from 40 cells in 5 independent experiments (control) and *n* = 238 bundles, pooled from 26 cells in 5 independent experiments (NuMA-KO+STLC). (F) Helicity of individual interpolar microtubule bundles, measured in degrees rotated (θ) around the pole-to-pole axis per µm traversed (d) along the pole-to-pole axis for each bundle. Schematic illustration of the helicity measurement shown in inset. Plot includes the same bundles tracked in (E). ****, p < 0.00005, two-sample t-test. Lines represent mean ± s.d. See also Figure S3. (G) Representative confocal images of control and NuMA-KO+STLC RPE1 anaphase cells stably expressing GFP-tubulin (not shown) and mCherry-H2B (cyan, maximum intensity projections, single frame from live imaging), showing lagging chromosomes in the NuMA-KO+STLC cell. Scale bars = 5 µm. (H) Percentage of anaphase cells with lagging chromosomes or chromosome bridges, showing increased segregation defects in NuMA-KO+STLC cells. *n* = 84 control cells pooled from 6 independent experiments; *n* = 35 NuMA-KO+STLC cells pooled from 5 independent experiments. ****, p < 0.00005, Fisher’s exact test.

Unexpectedly, we observed that in contrast with control spindles, doubly inhibited spindles were highly twisted in anaphase. Interpolar microtubule bundles followed a left-handed helical path around the spindle (Figure 4D). While doubly inhibited spindles exhibited twist to a small degree at metaphase (Figures S3A and S3B), the phenotype was much more pronounced and consistently left-handed after anaphase onset. To quantify this effect, we imaged z-stacks of anaphase spindles and tracked interpolar microtubule bundles in three-dimensional space. Viewing these trajectories along the pole-to-pole axis, interpolar bundles in doubly inhibited spindles had a helicity of −6.3 ± 3.4 °/µm, a 17-fold increase over control bundles’ helicity of −0.4 ± 2.0 °/µm (mean ± s.d.; Figures 4E, 4F, and S3C; Movie S6). Mean helicity was not correlated with anaphase spindle length (*r* = −0.04, Figure S3D), suggesting that spindle twist does not markedly increase or decrease as anaphase progresses. Together, these findings reveal an unexpected role for the opposing motor activities of NuMA/dynein and Eg5: although not required for linear force balance in the pole-to-pole axis (Figure 1F), they are required for rotational force balance in the anaphase spindle.

Finally, we asked whether chromosome segregation fidelity was preserved in doubly inhibited spindles. The incidence of chromosome segregation errors—defined here as lagging chromosomes or chromosome bridges—was significantly higher in NuMA-KO+STLC spindles than in controls (45.7% vs 7.1%; Figures 4G and 4H; Movie S5). Thus, while NuMA and Eg5 are not required for efficient spindle elongation, they are instead required at anaphase for the spindle’s straight long-range architecture and for accurate chromosome segregation.

## Discussion

The conserved presence of opposing extensile and contractile force-generators, despite their expendability for bipolar spindle formation, presents a long-standing paradox in spindle assembly. Our approach of inhibiting Eg5 and deleting NuMA, abrogating dynein’s role in minus end clustering but not its roles at the kinetochore, allows us to probe the roles of this opposing motor activity while preserving metaphase attachments and permitting anaphase progression. Here, we show that although NuMA- and Eg5-doubly inhibited human spindles appear strikingly similar to controls, they fail in key ways. We find that they are mechanically fragile at metaphase as well as dramatically twisted and error-prone at anaphase, defects that we propose stem from their altered dynamics, organization, and material properties (Figure 5). Thus, while partially redundant motors can establish spindle shape and support anaphase progression, the opposing activities of NuMA/dynein and Eg5 are required to build a spindle that can maintain its structure and accurate function despite internal and external pushes, pulls, and torques.

**Figure 5.**
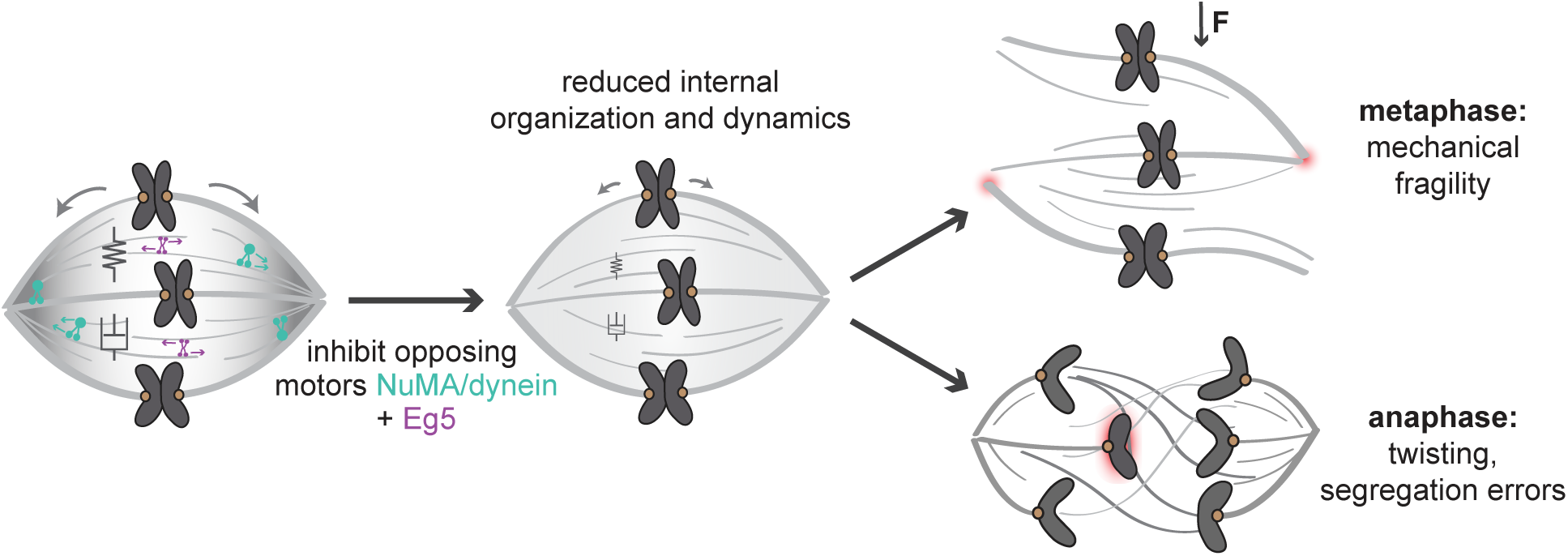
Model for Opposing Active Stresses Providing Mechanical and Functional Robustness to the Human Spindle. The spindle has opposing contractile and extensile stresses generated by NuMA/dynein (green) and Eg5 (purple), respectively (left). Without these opposing active stresses (center), the human spindle retains its steady-state shape and size and undergoes anaphase, but has reduced internal organization (gray gradient) and dynamics (gray arrows). These spindles are more structurally fragile when subjected to force (F vector) at metaphase (top right; red shading denotes pole fragmentation), become highly twisted at anaphase, and exhibit chromosome segregation errors (lower right; red shading denotes lagging chromosome). We propose that opposing active stresses give rise to mechanical and functional robustness by increasing the spindle’s organization and dynamics, and by tuning its material properties (springs, elasticity; dashpots, viscosity) to limit the magnitude and timescale of allowed deformations. Together, this work suggests a design principle whereby opposing active force generators promote mechanical and functional robustness of cellular machines.

We show that beyond their contributions to global spindle shape, NuMA and Eg5 are together essential to establishing the spindle’s locally specialized microtubule organization and dynamics (Figure 2). This raises the question of through which mechanisms they do so. The homogenized tubulin intensity distribution in doubly inhibited spindles could reflect deregulated microtubule transport or length (Brugues et al., 2012), or could result from aberrant microtubule nucleation, potentially through Eg5’s regulation of the nucleation factor TPX2 (Ma et al., 2010). The slower flux in doubly inhibited spindles could result from reduced NuMA/dynein- and Eg5-mediated poleward transport of non-kinetochore microtubules (Lecland and Luders, 2014) and crosslinking of these microtubules to k-fibers (Steblyanko et al., 2020). Notably, while one proposed role of flux is spindle length regulation (Fu et al., 2015; Steblyanko et al., 2020), the decoupling of flux and spindle length we observe in doubly inhibited spindles is not easily consistent with this idea. NuMA and Eg5’s roles in spindle organization and flux may instead have consequences for the spindle’s mechanical integrity and functional accuracy.

We find that spindles are more mechanically fragile without opposing NuMA/dynein and Eg5 activity (Figure 3). Cell confinement revealed that doubly inhibited spindles are more compliant and less able to maintain a steady-state structure under force. This could stem from lower microtubule enrichment at poles (Figure 2B) (Takagi et al., 2019), a less dynamic spindle (Figure 2E), reduced passive crosslinking, or decreased active stresses throughout the spindle. All of these are ways in which reduced opposing motor activity could impair the spindle’s ability to distribute and dissipate force, and thereby change the magnitude and timescale of the spindle’s deformation under force. Motors broadly regulate the material properties of microtubule networks, such as their elasticity and viscosity (Brugues and Needleman, 2014; Oriola et al., 2020; Shimamoto et al., 2011). Looking forward, combining the experimental system used here with approaches such as microneedle manipulation (Gatlin et al., 2010; Shimamoto et al., 2011; Suresh et al., 2020; Takagi et al., 2019) will enable us to understand how opposing active motors quantitatively tune the spindle’s emergent mechanical properties.

Our anaphase observations indicate that spindles have structural as well as functional defects without NuMA and Eg5. They reliably progress to anaphase and efficiently elongate, supporting a model where Eg5-independent sliding within the spindle can generate the bulk of the force required for chromosome segregation (Vukusic et al., 2017; Vukusic et al., 2019; Yu et al., 2019). However, in contrast to controls, doubly inhibited anaphase spindles exhibit strong left-handed twist, suggesting that they have an imbalance in torques. Multiple mitotic motors have an intrinsic chirality to their stepping motion *in vitro* (Can et al., 2014; Mitra et al., 2018; Nitzsche et al., 2016; Yajima et al., 2008), which can twist microtubules around each other (Mitra et al., 2020). At the cellular scale, left-handed helicity of a smaller magnitude (approximately −2°/µm) exists in metaphase and anaphase human spindles, but this twist is reduced by Eg5 inhibition (Novak et al., 2018; Trupinic et al., 2020). Thus, as Eg5 is inhibited in the anaphase spindles probed here, a different mechanism must produce the left-handed torque. One possibility is that spindles lacking NuMA/dynein and Eg5 activity are twisted due to abnormally high torques generated by the motors that compensate for their absence. However, because doubly inhibited spindles are more mechanically deformable (Figure 3) and because they over-elongate in anaphase (Figure 4B), we favor a model in which they are instead more torsionally compliant. Regardless of its molecular origin, the appearance of twist upon inhibition of NuMA and Eg5 raises the question of how the cell builds a micron-scale, near-achiral spindle from nanometer-scale chiral events. This requires a balancing of three-dimensional rotational forces over large length scales, through mechanisms that remain poorly understood. The doubly inhibited spindles we generate here may provide a system to uncover these mechanisms, and to address the functional impact of twist in the anaphase spindle. While the chromosome segregation errors we observe could arise from the re-clustering of microtubules in turbulent spindles (Ganem et al., 2009) or from altered microtubule dynamics (Bakhoum et al., 2009) or flux (Ganem et al., 2005; Matos et al., 2009), they could also arise from the spindle’s anaphase twist. For example, segregating chromosomes might follow more complex, entangled trajectories, or the elongating spindle could generate an increased non-productive force component.

Overall, our findings indicate that the opposing activities of NuMA/dynein and Eg5 are critical for the spindle’s mechanical and functional robustness, allowing the spindle to withstand force and accurately segregate chromosomes despite its dynamic molecular parts. An energy-accuracy tradeoff has been demonstrated experimentally and theoretically in biochemical networks: for instance, repeated energy-consuming cycles of kinase and phosphatase activity synchronize cell cycle timing in zebrafish embryos (Rodenfels et al., 2019), and phase coherence scales with energy dissipation in a variety of biochemical oscillators (Cao et al., 2015). We propose that opposing spindle motors provide a mechanical analog, where the spindle’s structural integrity and functional accuracy incur an energetic cost beyond that required to establish spindle structure. Opposing active force generators may constitute a physical design principle that underlies robustness in other dynamic, self-organizing cellular structures, such as cell-cell junctions.

## Acknowledgements

We thank Christina Hueschen and Andrea Serra-Marques for inducible RPE1 NuMA-KO and DHC-KO cell lines, Iain Cheeseman and Kara McKinley for reagents, and Dan Needleman, Margaret Gardel, and Kendra Burbank for helpful discussions. We thank Christina Hueschen, Rob Phillips, and members of the Dumont lab for critical reading of the manuscript. Photomarking experiments were performed at the UCSF Center for Advanced Light Microscopy – Nikon Imaging Center with assistance from Kari Herrington and SoYeon Kim, on instruments obtained using funding from the NIH (5R35GM118119), the UCSF Program in Breakthrough Biomedical Research, the Sandler Foundation, the UCSF Research Resource Fund Award, and the Howard Hughes Medical Institute. This work was supported by NIH DP2GM119177, NIH R01GM134132, NIH R35GM136420, NSF CAREER 1554139, and NSF 1548297 Center for Cellular Construction (S.D.); the Chan Zuckerberg Biohub, Rita Allen Foundation and Searle Scholars’ Program (S.D.), and an NSF Graduate Research Fellowship, Fannie and John Hertz Foundation Fellowship, and William K. Bowes Jr. & Ute Bowes Discovery Fellowship (L.N.).

## Author Contributions

Conceptualization, L.N. and S.D.; Methodology, L.N., N.H.C. and S.D.; Software, L.N.; Validation, L.N. and N.H.C.; Formal Analysis, L.N.; Investigation, L.N. and N.H.C.; Resources, L.N., N.H.C. and S.D.; Data Curation, L.N.; Writing – Original Draft, L.N.; Writing – Review & Editing, L.N., N.H.C. and S.D.; Visualization, L.N.; Supervision, L.N. and S.D.; Funding Acquisition, L.N. and S.D.

## Methods

### RESOURCE AVAILABILITY

#### Lead Contact

Further information and requests for resources and reagents should be directed to and will be fulfilled by the Lead Contact, Sophie Dumont.

#### Materials Availability

All unique cell lines generated in this study are available from the Lead Contact without restriction.

#### Data and Code Availability

Datasets and code generated for this study are available from the Lead Contact without restriction.

### EXPERIMENTAL MODEL AND SUBJECT DETAILS

All cell lines were generated from an hTERT-RPE1 cell line (female human retinal epithelial cells) stably expressing neomycin-resistant tet-on SpCas9, a gift from I. Cheeseman (McKinley and Cheeseman, 2017). Cell lines additionally expressed a puromycin-selectable sgRNA targeting NuMA or dynein heavy chain (Hueschen et al., 2019). All cell lines were cultured at 37° and 5% CO_2_ in DMEM/F12 (11320, Thermo Fisher) supplemented with 10% tetracycline-screened FBS (PS-FB2, Peak Serum). Fluorescently tagged proteins were introduced by transduction with blasticidin-resistant GFP-tubulin, mCherry-H2B, or PA-GFP-tubulin lentivirus, produced in HEK293T cells, supplemented with 10 µg/ml polybrene. Cell lines were selected with 5 µg/ml puromycin and 5 µg/ml blasticidin. SpCas9 expression was induced by the addition of 1 µg/ml doxycycline hyclate 4 days before each experiment, refreshed after 24 and 48 h.

### METHOD DETAILS

#### Transfection and Small Molecule Treatments

For siRNA knockdowns, cells were transfected with 50 pmol siRNA targeting luciferase as a negative control (5’-CGUACGCGGAAUACUUCGA-3’, 48 h), HSET (5’-UCAGAAGCAGCCCUGUCAA-3’, 48 h) (Cai et al., 2009), or Kif15 (5’-GGACAUAAAUUGCAAAUAC-3’, 24 h) (Vanneste et al., 2009) using Lipofectamine RNAiMAX (13778075, Thermo Fisher) according to the manufacturer’s recommendations. Chromosomes were labeled in the inducible DHC-KO cell line (Figures 1C, 1I, 1J) by incubating cells in 1 µM SiR-DNA and 10 µM verapamil (CY-SC007, Cytoskeleton Inc.) for 60 min prior to imaging. For photomarking experiments (Figures 2D and 2E), microtubules were labeled by incubating cells with 100 nM SiR-tubulin and 10 µM verapamil (CY-SC002, Cytoskeleton Inc.) for 60 min prior to imaging. For all experiments, cells were synchronized at the G2/M checkpoint by overnight treatment with 9 µM of the Cdk1 inhibitor RO-3306. Cells were released into mitosis by 4 washes in warm media, after which cells were imaged from prometaphase (controls, approximately 30 min after washout) or from reaching the turbulent state (NuMA- or DHC-KO, approximately 60 min after washout). Eg5 motor activity was inhibited by addition of S-trityl-L-cysteine (final concentration 5 µM, 164739, Sigma). To bypass the spindle assembly checkpoint (Figure 1J), the MPS1 inhibitor reversine (final concentration 500 nM, R3904, Sigma) was added 45 min after STLC. To disrupt F-actin (Figure 1K), Latrunculin A (final concentration 500 nM, L12370, Invitrogen) was added at the same time as STLC.

#### Microscopy

For live imaging, cells were plated onto #1.5 glass-bottom 35 mm dishes coated with poly-D-lysine (P35G-1.5-20-C, MatTek Life Sciences) and imaged in a humidified stage-top incubator maintained at 37° and 5% CO_2_ (Tokai Hit). Fixed and live cells were imaged on a spinning disk (CSU-X1, Yokogawa) confocal inverted microscope (Eclipse Ti-E, Nikon Instruments) with the following components: Di01-T405/488/561/647 head dichroic (Semrock); 405 nm (100 mW), 488 nm (150 mW), 561 nm (100 mW) and 642 nm (100 mW) diode lasers; ET455/50M, ET525/50M, ET630/75M, and ET705/72M emission filters (Chroma Technology); and a Zyla 4.2 sCMOS camera (Andor Technology). Images were acquired with a 100× 1.45 Ph3 oil objective using MetaMorph 7.10.3.279 (Molecular Devices). Photomarking experiments (Figures 2D and 2E) were performed on an OMX-SR inverted microscope (GE Healthcare) with the following components: three PCO Edge 5.5 sCMOS cameras; an environmental chamber maintained at 37° and 5% CO_2_ (GE Healthcare); and a Plan ApoN 60× 1.42 oil objective. Photoactivation was performed with a single 20 ms pulse of 405 nm light targeted to a rectangular region of interest.

#### Immunofluorescence

For immunofluorescence, cells were plated onto acid-cleaned #1.5 25 mm coverslips coated with 0.1% gelatin solution. Cells were fixed in methanol at −20°C for 3 min, washed with TBST (0.05% Triton-X-100 in TBS), and blocked with 2% BSA in TBST. Antibodies were diluted in TBST + 2% BSA and incubated overnight at 4°C (primary antibodies) or 45 min at room temperature (secondary antibodies). DNA was labeled with 1 µg/ml Hoechst 33342 for 20 min, prior to mounting on slides with ProLong Gold Antifade Mountant (P36934, Thermo Fisher). The following primary antibodies were used: mouse anti-α-tubulin (1:1,000, T6199, Sigma), rat anti-α-tubulin (1:500, MCA77G, Bio-Rad), rabbit anti-NuMA (1:300, NB500-174, Novus Biologicals), and mouse anti-α-tubulin AlexaFluor 488 conjugate (1:50, added with secondary antibodies, 8058S, Cell Signaling Technology). The following secondary antibodies were used at a 1:400 dilution: goat anti-rabbit AlexaFluor 568 and AlexaFluor 647 (A-11011 and A-21244, Thermo Fisher), goat anti-rat AlexaFluor 488 (A-11006, Thermo Fisher), and goat anti-mouse AlexaFluor 488 (A-11001, Thermo Fisher). Brightness/contrast for each channel was scaled identically within each immunofluorescence experiment shown.

#### Western Blotting

Cells in 6-well plates were lysed, and protein extracts were collected after centrifugation at 4°C for 30 min. Protein concentrations were measured using a Bradford assay kit (Bio-Rad), and equal concentrations of each sample were separated on a 3-8% Tris-Acetate or 4-12% Bis-Tris gel (Invitrogen) by SDS-PAGE and transferred to a nitrocellulose membrane. Membranes were blocked with 4% milk, incubated in primary antibodies overnight at 4°C, and incubated with HRP-conjugated secondary antibodies for 1 h. Proteins were detected using SuperSignal West Pico or Femto chemiluminescent substrates (Thermo Fisher). The following primary antibodies were used: mouse anti-α-tubulin (1:5,000, T6199, Sigma), rabbit anti-NuMA (1:1,000, NB500-174, Novus Biologicals), rabbit anti-Kif15 (1:500, A302-706A, Bethyl Laboratories), and mouse anti-KifC1 (M-63; 1:500, sc-100947, Santa Cruz Biotechnology). The following secondary antibodies were used at a 1:10,000 dilution: goat anti-mouse IgG-HRP (sc-2005, Santa Cruz Biotechnology) and mouse anti-rabbit IgG-HRP (sc-2357, Santa Cruz Biotechnology).

#### Cell Confinement

Cells were confined as described previously (Guild et al., 2017), using a suction cup device adapted from Le Berre et al. (2012). Briefly, PDMS pillars 5 µm in height (200 µm diameter, 700 µm spacing) were attached to a 10 mm-diameter coverslip, and were lowered onto cells using negative pressure generated manually using a 1 ml syringe. Pillars were gradually lowered onto cells over ∼2 min, and maximum confinement (at a cell height of 5 µm) was sustained for an additional 20 min. Cells were excluded from analysis if the final confined height was >5 µm, suggesting that the cell’s surroundings on the coverslip prevented full confinement, or if the separation between sister chromosomes became indistinguishable, suggesting chromosome decondensation, e.g. resulting from cell rupture.

### QUANTIFICATION AND STATISTICAL ANALYSIS

#### Quantification of Spindle Shape and Failure

Spindle length and width were measured manually using the line selection tool in FIJI (ImageJ version 2.0.0-rc-69/1.52p). For control and NuMA-KO+STLC cells, length was measured as the distance between the two spindle poles, and width was measured at the widest part of the spindle across the metaphase plate. Aspect ratio was determined by dividing length by width. For turbulent NuMA-KO spindles and compressed spindles after structural failure, spindle axis directions were approximated from chromosome positions. For Figures 1F-1H, spindle dimensions were measured after reaching a bipolar metaphase (control and NuMA-KO+STLC) or 45 min after the start of imaging (NuMA-KO). For Figures 1I-1K, global spindle shape and anaphase entry were scored at 90 min after STLC addition, and cells that were imaged for <90 min were excluded. Spindle failure (Figure 3F) was defined as a loss of visible connectivity between k-fibers and the pole.

#### Quantification of NuMA Levels via Immunofluorescence

To compare NuMA intensity in control cells versus cells in which NuMA knockout had been induced, we quantified sum intensity projections of 21 z-planes spaced 0.35 µm apart. Using a custom MATLAB program (version R2020a), cell areas were segmented using a low tubulin threshold, and mean NuMA and tubulin intensities were measured within this region. NuMA intensities were normalized for each cell by dividing by the corresponding tubulin intensity. For the +DOX condition, only spindles with a disorganized phenotype (a single snapshot of a turbulent spindle) were analyzed, consistent with the criterion of spindle turbulence used for all live imaging experiments.

#### Fluorescence Intensity Profiles

Fluorescence intensity profiles along the pole-to-pole axis (Figures 2B and 2C) were quantified from sum intensity projections of 21 z-planes spaced 0.35 µm apart. Using a custom MATLAB program, images of tubulin fluorescence were passed through a median filter (3×3 pixels) and spindle areas were segmented using a tubulin intensity threshold. Based on the major axis angle of the segmented spindle, images were rotated so that the pole-to-pole axis was horizontal. At each of 21 positions (0%, 5%, 10%…100%) along the pole-to-pole axis, the mean tubulin and Hoechst intensities were calculated from the 1-pixel-wide column of all pixels contained within the spindle boundaries. Finally, these 21-point profiles were normalized to the maximum value for each spindle.

#### Flux Rate

SiR-tubulin image sequences were aligned using a Rigid Body transformation, and the corresponding PA-GFP-tubulin image sequence was registered using the MultiStackReg plugin (version 1.45) to remove overall spindle drift. FIJI’s segmented line selection tool with spline fitting was used to trace 2-3 k-fibers per spindle, and kymographs were generated from the PA-GFP-tubulin channel for each k-fiber using the Multi Kymograph plugin with a linewidth of 3 pixels. In MATLAB, the intensity values in each kymograph were smoothed with a moving mean calculated over a sliding 5-pixel window, and the position of maximum intensity was determined for each timepoint. Linear regression was performed on the positions of these maxima to determine the rate that the photomark moved polewards, using the MATLAB fit function of type ‘poly1’.

#### Time Correlation Function of Spindle Shape

Timelapse image sequences were registered in FIJI using the Rigid Body option of the StackReg plugin (Thévenaz et al., 1998). Spindles were segmented in FIJI by smoothing, despeckling, background subtraction, and thresholding with Otsu’s method. In MATLAB, thresholded binary image sequences were cropped to a 33×33 µm box centered at the spindle’s centroid, and spindle masks were further refined by filling holes and removing small objects. The correlation coefficient was calculated, using the MATLAB corr2 function, between all pairs of binarized frames separated by lag time Δt, where Δt = 0.5, 1, 1.5, … 9.5 min. To determine shape correlation as a function of lag time, correlation coefficients were averaged for each lag time and fit to the exponential function 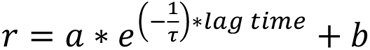 using MATLAB’s curve fitting tool (Hueschen et al., 2019).

#### Anaphase Segregation Rates

Cells analyzed in Figures 4B, 4C, and S2 were imaged every 30 s from late metaphase through telophase. Anaphase onset was defined as the first frame with detectable chromosome separation. In each frame, the distance between the two spindle poles and the distance between the centers of the two chromosome masses were measured manually with the line selection tool in FIJI. Elongation and segregation rates were determined by linear regression of data between t = 30 s and t = 180 s, using the MATLAB fit function of type ‘poly1’.

#### Microtubule Bundle Helicity

Helicity was analyzed similarly to the method described in Novak et al. (2018). We acquired z-stacks of GFP-tubulin-labeled spindles from live metaphase and anaphase cells. Z-axis calibration was performed using a FocalCheck slide #1 (F36909, Thermo Fisher), and the preservation of handedness throughout the optical train was validated by imaging a 3mm-diameter spring of known handedness with a 10× objective. Z-stacks were manually rotated in FIJI such that the pole-to-pole axis was horizontal. Image coordinates (x, y, z) were permuted to (z, x, y) in MATLAB, creating a series of spindle cross-sections as if viewed end-on from the pole (Movie S6). The rotated image stacks were background-subtracted and despeckled to facilitate bundle tracking. Spindle poles were marked and individual bundles were traced in FIJI using the MTrackJ plugin (Meijering et al., 2012), with cursor snapping to the bright centroid of a 15×15 pixel box enabled. In MATLAB, tracked bundle and pole positions were transformed so that both poles lay on the x-axis, accounting for spindle tilt. Tracked points were excluded if they lay outside the central 30-70% of the pole-to-pole axis. Bundles were excluded from further analysis if their mean radial distance from the central pole-to-pole axis was <2 µm, or if they contained fewer than 20 points (corresponding to a minimum track length of 1.16 µm). The angle between the first and last point in each bundle track was calculated with respect to the central pole-to-pole axis, and this angle was divided by the distance traversed along the pole-to-pole axis to calculate helicity.

#### Quantification of Chromosome Segregation Errors

Segregation errors (Figures 4G and 4H) were determined from z-stacks of mCherry-H2B fluorescence, acquired with 1 µm spacing and covering the entire spindle height at a single timepoint during live imaging. Segregation errors included lagging chromosomes, defined here as one or more chromosomes completely separated from the rest of the chromosome mass, and chromosome bridges, defined here as an extended chromosome pair connecting the two segregating masses.

#### Statistical Analysis

Details of statistical tests and sample sizes (number of cells and number of independent experiments) are provided in figure legends. Fisher’s exact tests were performed to compare categorical datasets, using the fishertest function in MATLAB for 2×2 comparisons and the fisher.test function in R (version 4.0.1) for 2×3 comparisons. Two-sided two-sample t-tests were performed to compare continuous datasets using the ttest2 function in MATLAB, based on the assumption that spindle length and width, flux rate, anaphase segregation rate, and helicity are approximately normally distributed. We used p < 0.05 as the threshold for statistical significance. Linear regressions (Figures 2E, 4B, 4C, S2A, and S2B) and exponential decay fits (Figure 3E) were performed in MATLAB.

## Supplemental Information

**Figure S1.**
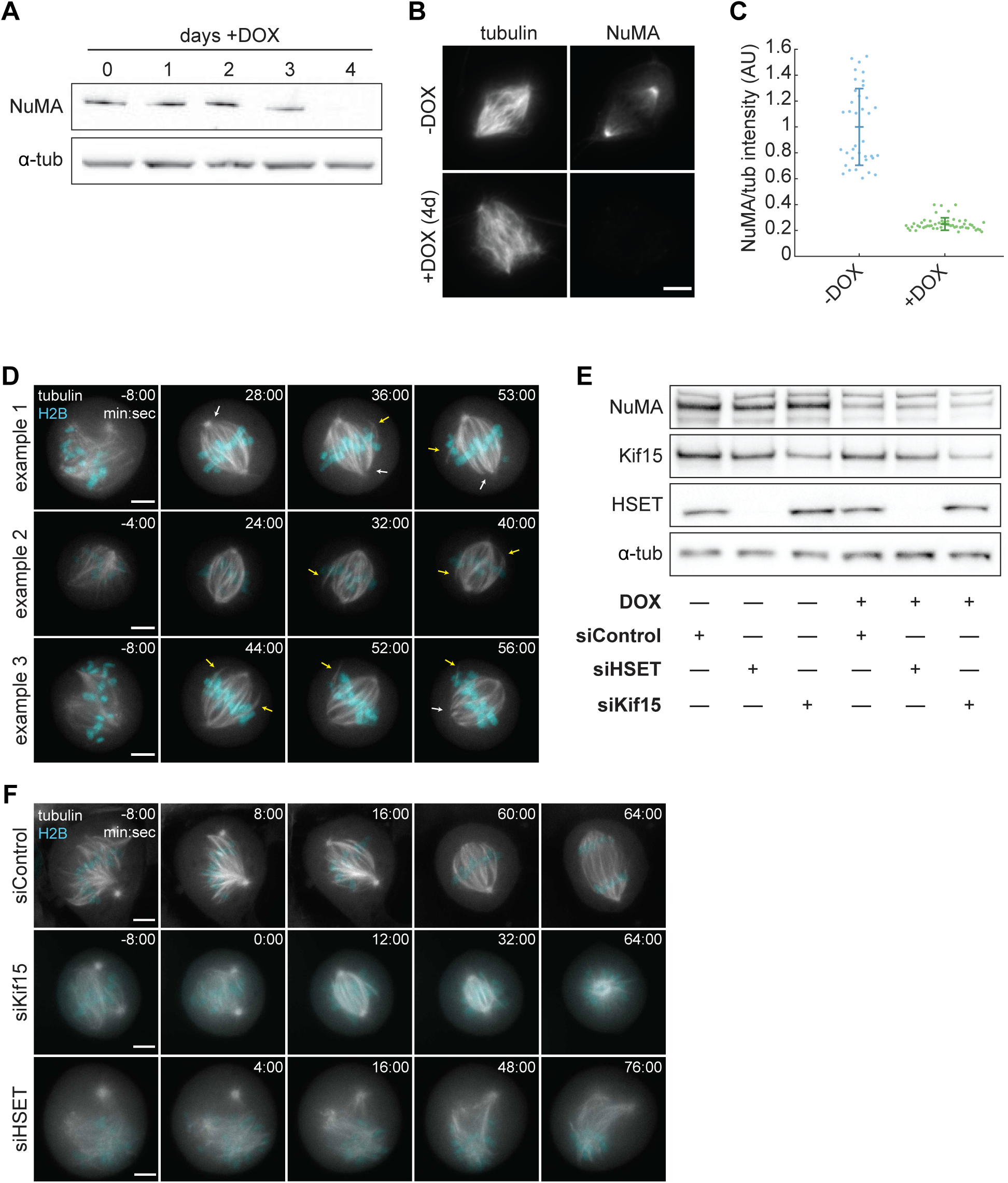
Additional Analysis of NuMA- and Eg5-Doubly Inhibited Spindles. (A) Western blot of NuMA levels in RPE1 cells after DOX-induced Cas9 expression for the indicated number of days. Tubulin is shown as a loading control. (B) Representative immunofluorescence images (sum projections) of inducible NuMA-KO RPE1 cells with or without DOX induction of Cas9 for 4 days. Cells are stained for tubulin (left) and NuMA (right). Scale bar = 5 µm. (C) Quantification of NuMA intensity in immunofluorescence images (sum projections), normalized to tubulin intensity. Control (-DOX) metaphase cells were compared to NuMA-KO (+DOX) cells with turbulent spindles. n = 39 cells (-DOX) and 53 cells (+DOX) from one experiment. Lines represent mean ± s.d. (D) Timelapse confocal images of RPE1 NuMA-KO+STLC cells, stably expressing GFP-tubulin (gray) and mCherry-H2B (cyan). Spindles recover bipolarity after 5 µM STLC addition at time 0:00. White arrows indicate pole unfocusing, and yellow arrows indicate k-fibers that dynamically splay and reincorporate into the bipolar spindle. The tubulin channel shows maximum intensity projections of 5 planes spaced 0.5 µm apart, while the H2B channel shows single z-planes. Scale bars = 5 µm. (E) Western blot of NuMA, Kif15, and HSET levels in RPE1 cells with or without DOX induction of Cas9 for 4 days, and transfected with siRNA targeting luciferase (Control) or HSET for 48 hours, or siRNA targeting Kif15 for 24 hours. Tubulin is shown as a loading control. While NuMA-KO efficiency at the population level varied between experiments, NuMA-KO was verified in each individual cell, for all experiments, based on NuMA immunofluorescence or live imaging of the turbulent phenotype prior to STLC addition. (F) Timelapse confocal images of RPE1 NuMA-KO+STLC cells stably expressing GFP-tubulin (gray) and mCherry-H2B (cyan), transfected with siRNA targeting luciferase (Control), HSET, or Kif15. After 5 µM STLC addition at time 0:00, the siControl spindle recovers bipolarity, the siKif15 spindle collapses into a monopole, and the siHSET spindle remains turbulent. The tubulin channel shows maximum intensity projections of 5 planes spaced 0.5 µm apart, while the H2B channel shows single z-planes. Scale bars = 5 µm.

**Figure S2.**
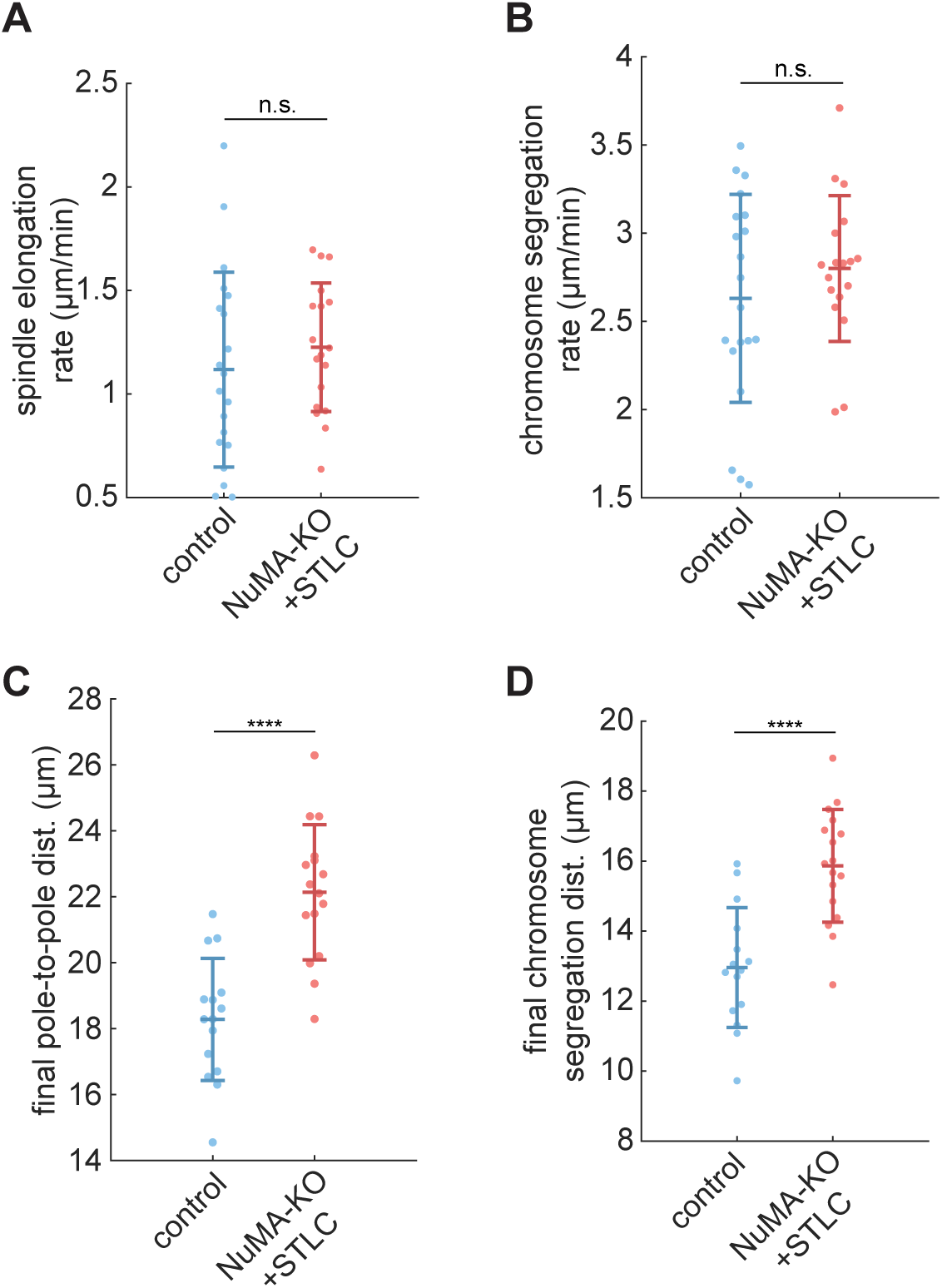
Comparison of Anaphase Movements in Control and Doubly Inhibited Spindles. (A) Average spindle elongation rate (change in pole-to-pole distance per min) in the period from 30-180 s after anaphase onset, corresponding to the gray shaded box in Figure 4B, in control and NuMA-KO+STLC RPE1 cells. (B) Average chromosome segregation rate (change in distance between chromosome masses per min) in the period from 30-180 s after anaphase onset, corresponding to the gray shaded box in Figure 4C, in control and NuMA-KO+STLC cells. (C) Pole-to-pole distance 9 min after anaphase onset, when spindle elongation had largely ceased, in control and NuMA-KO+STLC cells. (D) Chromosome segregation distance 9 min after anaphase onset, when chromosome segregation had largely ceased, in control and NuMA-KO+STLC cells. For (A)-(D), each dot represents one cell. Data include the same 20 control and 18 NuMA-KO+STLC cells shown in Figures 4B and 4C, pooled from 4 independent experiments. ****, p < 0.00005, n.s., not significant, two-sample t-test. Lines indicate mean ± s.d.

**Figure S3.**
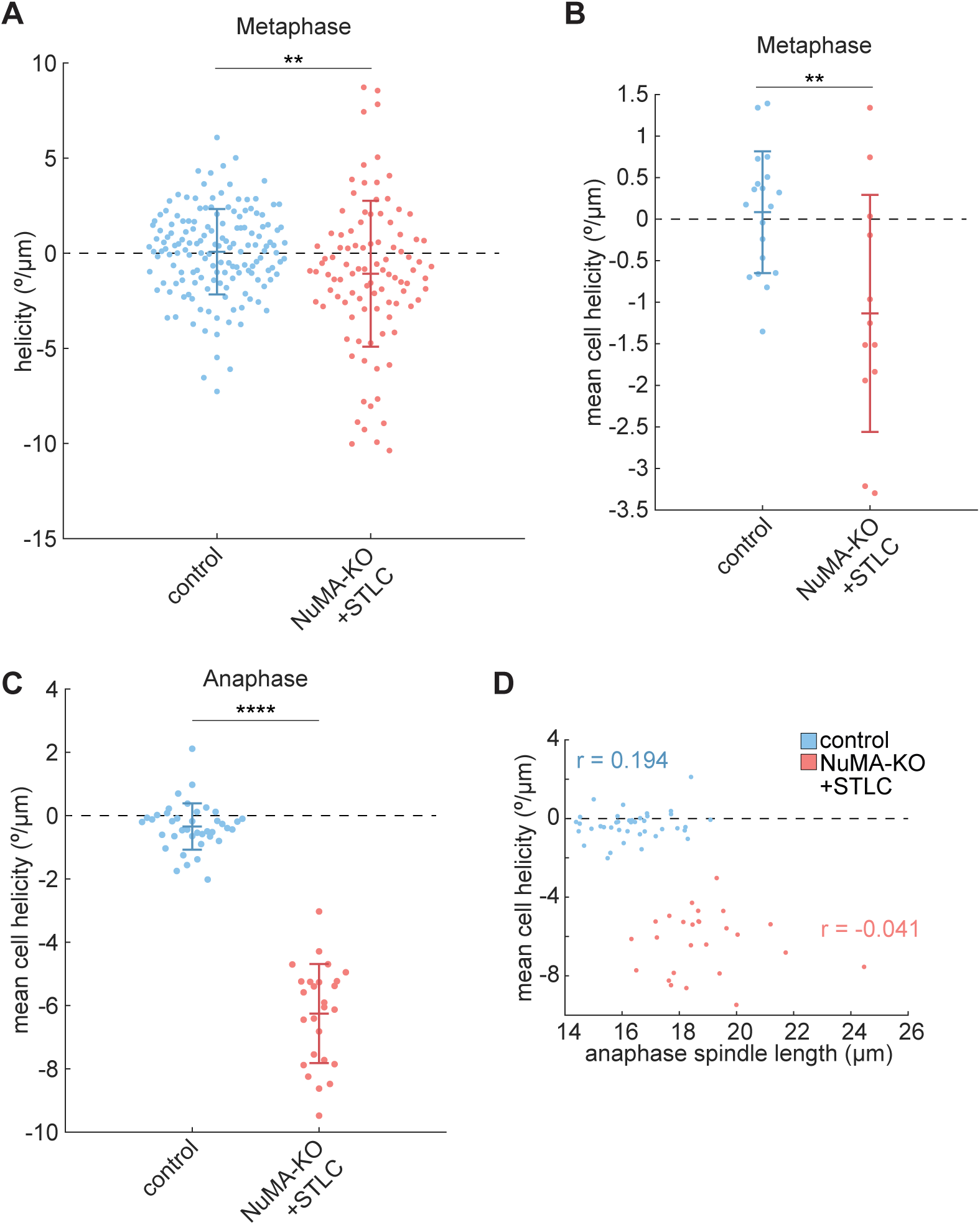
Additional Analysis of Spindle Twist in Control and Doubly Inhibited Spindles. (A) Helicity of individual metaphase k-fibers, measured in degrees rotated around the pole-to-pole axis per µm traversed along the pole-to-pole axis. Each dot represents one k-fiber. The mean helicity of control k-fibers is 0.08°/µm, while NuMA-KO+STLC k-fibers have a left-handed average helicity of −1.08°/µm. n = 155 k-fibers from 19 cells (control) and 99 k-fibers from 12 cells (NuMA-KO+STLC), pooled from 3 (control) or 2 (NuMA-KO+STLC) independent experiments. **, p = 0.0027, two-sample t-test. Lines indicate mean ± s.d. (B) Mean helicity of metaphase k-fibers per cell (each dot represents one cell), including the same data as (A). Metaphase twist was not consistently left- or right-handed for control or NuMA-KO+STLC spindles. **, p = 0.0039, two-sample t-test. Lines indicate mean ± s.d. (C) Mean helicity of anaphase interpolar bundles per cell (each dot represents one cell), including the same data as Figures 4E and 4F. NuMA-KO+STLC spindles all exhibit left-handed twist, while control spindle twist is not consistently left- or right-handed. n = 40 cells (control) and 26 cells (NuMA-KO+STLC), pooled from 5 independent experiments. ****, p < 0.00005. Lines indicate mean ± s.d. (D) Scatterplot of anaphase spindle length vs. mean helicity of interpolar bundles (each dot represents one cell), including the same data as (C) and Figures 4E and 4F. Mean helicity is not correlated with the extent of anaphase spindle elongation for either control or NuMA-KO+STLC cells, suggesting that spindles do not markedly twist or untwist as anaphase progresses.

**Movie S1. Turbulent DHC-KO and NuMA-KO Spindles Recover Bipolarity After Eg5 Inhibition**

Timelapse confocal imaging of a representative RPE1 DHC-KO cell (left) and a representative RPE1 NuMA-KO cell (right), recovering bipolarity after the addition of 5 µM STLC at time 00:00 (min:s). The NuMA-KO cell enters anaphase at 51:45, while the DHC-KO cell does not enter anaphase within the observation period. Both cells stably express GFP-tubulin (gray; maximum intensity projections of 5 planes spaced 0.5 µm apart). Chromosomes (cyan; single z-plane) are labeled with SiR-DNA (DHC-KO, left) or stably expressed mCherry-H2B (NuMA-KO, right). Movie corresponds to still images in Figures 1C and 1D.

**Movie S2. HSET and Kif15 Provide Partially Redundant Contractile and Extensile Activity to Establish Bipolarity in Doubly Inhibited Spindles**

Timelapse confocal imaging of representative RPE1 NuMA-KO cells transfected with siRNA targeting luciferase (control), Kif15, or HSET, stably expressing GFP-tubulin (gray; maximum intensity projection of 5 planes spaced 0.5 µm apart) and mCherry-H2B (cyan; single plane). After 5 µM STLC addition at time 00:00:00 (h:min:s), the control cell establishes a bipolar spindle, the cell depleted of Kif15 establishes a monopolar spindle, and the cell depleted of HSET remains turbulent. Movie corresponds to still images in Figure S1F.

**Movie S3. Doubly Inhibited Spindles Exhibit Slower Microtubule Flux**

Timelapse widefield imaging of representative RPE1 control and NuMA-KO+STLC cells, stably expressing PA-GFP-tubulin (green) and labeled with 100 nM SiR-tubulin (gray). The PA-GFP-tubulin channel alone is shown below, with asterisks marking spindle poles. A stripe of tubulin fluorescence is photoactivated near the spindle equator at time 00:00 (min:s), after which the photomark translocates towards the pole more rapidly in the control cell. Movie corresponds to still images in Figure 2D.

**Movie S4. Doubly Inhibited Spindles Fracture More Readily During Confinement**

Timelapse confocal imaging of representative RPE1 control and NuMA-KO+STLC cells, stably expressing GFP-tubulin (gray) and mCherry-H2B (cyan), under confinement beginning at time 00:00 (min:s). Confinement is applied during the first 2 minutes and is sustained for 20 min. Both spindles widen and lengthen, but the NuMA-KO+STLC spindle loses its structural integrity while the control spindle remains intact. Movie corresponds to still images in Figure 3B.

**Movie S5. Doubly Inhibited Spindles Segregate Chromosomes to Greater Distances and with More Errors**

Timelapse confocal imaging of representative RPE1 control and NuMA-KO+STLC cells, stably expressing GFP-tubulin (gray) and mCherry-H2B (cyan), after anaphase onset at time 00:00 (min:s). The spindle elongates more, and chromosomes segregate farther, in the doubly inhibited cell. Arrow denotes a lagging chromosome. Movie corresponds to still images in Figure 4A.

**Movie S6. End-on Views of Control and Doubly Inhibited Anaphase Spindles, Revealing Left-Handed Twist**

Single-timepoint z-stacks of live-imaged anaphase spindles, after rotation of the coordinate system to view each spindle along its pole-to-pole axis. Representative examples of RPE1 control (left) and NuMA-KO+STLC (right) cells, stably expressing GFP-tubulin (gray) and mCherry-H2B (not shown). Interpolar microtubule bundles in control spindles follow straight trajectories. In contrast, interpolar bundles in doubly inhibited spindles follow helical trajectories, with each bundle tracing a clockwise arc as it approaches the viewer, revealing left-handed twist.

## References

Aist, J.R., Liang, H., and Berns, M.W. (1993). Astral and spindle forces in PtK2 cells during anaphase B: a laser microbeam study. J Cell Sci 104 (Pt 4), 1207–1216.

Bakhoum, S.F., Genovese, G., and Compton, D.A. (2009). Deviant kinetochore microtubule dynamics underlie chromosomal instability. Curr Biol 19, 1937–1942.

Blangy, A., Lane, H.A., d’Herin, P., Harper, M., Kress, M., and Nigg, E.A. (1995). Phosphorylation by p34cdc2 regulates spindle association of human Eg5, a kinesin-related motor essential for bipolar spindle formation in vivo. Cell 83, 1159–1169.

Brugues, J., and Needleman, D. (2014). Physical basis of spindle self-organization. Proc Natl Acad Sci U S A 111, 18496–18500.

Brugues, J., Nuzzo, V., Mazur, E., and Needleman, D.J. (2012). Nucleation and transport organize microtubules in metaphase spindles. Cell 149, 554–564.

Cai, S., Weaver, L.N., Ems-McClung, S.C., and Walczak, C.E. (2009). Kinesin-14 family proteins HSET/XCTK2 control spindle length by cross-linking and sliding microtubules. Mol Biol Cell 20, 1348–1359.

Cameron, L.A., Yang, G., Cimini, D., Canman, J.C., Kisurina-Evgenieva, O., Khodjakov, A., Danuser, G., and Salmon, E.D. (2006). Kinesin 5-independent poleward flux of kinetochore microtubules in PtK1 cells. J Cell Biol 173, 173–179.

Can, S., Dewitt, M.A., and Yildiz, A. (2014). Bidirectional helical motility of cytoplasmic dynein around microtubules. eLife 3, e03205.

Cao, Y., Wang, H., Ouyang, Q., and Tu, Y. (2015). The free energy cost of accurate biochemical oscillations. Nat Phys 11, 772–778.

Crowder, M.E., Strzelecka, M., Wilbur, J.D., Good, M.C., von Dassow, G., and Heald, R. (2015). A comparative analysis of spindle morphometrics across metazoans. Curr Biol 25, 1542–1550.

Dumont, S., and Mitchison, T.J. (2009). Compression regulates mitotic spindle length by a mechanochemical switch at the poles. Curr Biol 19, 1086–1095.

Elting, M.W., Suresh, P., and Dumont, S. (2018). The Spindle: Integrating Architecture and Mechanics across Scales. Trends Cell Biol 28, 896–910.

Ferenz, N.P., Paul, R., Fagerstrom, C., Mogilner, A., and Wadsworth, P. (2009). Dynein antagonizes Eg5 by crosslinking and sliding antiparallel microtubules. Curr Biol 19, 1833–1838.

Florian, S., and Mayer, T.U. (2012). The functional antagonism between Eg5 and dynein in spindle bipolarization is not compatible with a simple push-pull model. Cell Rep 1, 408–416.

Foster, P.J., Furthauer, S., Shelley, M.J., and Needleman, D.J. (2015). Active contraction of microtubule networks. eLife 4.

Fu, J., Bian, M., Xin, G., Deng, Z., Luo, J., Guo, X., Chen, H., Wang, Y., Jiang, Q., and Zhang, C. (2015). TPX2 phosphorylation maintains metaphase spindle length by regulating microtubule flux. J Cell Biol 210, 373–383.

Gaglio, T., Saredi, A., Bingham, J.B., Hasbani, M.J., Gill, S.R., Schroer, T.A., and Compton, D.A. (1996). Opposing motor activities are required for the organization of the mammalian mitotic spindle pole. J Cell Biol 135, 399–414.

Ganem, N.J., Godinho, S.A., and Pellman, D. (2009). A mechanism linking extra centrosomes to chromosomal instability. Nature 460, 278–282.

Ganem, N.J., Upton, K., and Compton, D.A. (2005). Efficient mitosis in human cells lacking poleward microtubule flux. Curr Biol 15, 1827–1832.

Gassmann, R., Holland, A.J., Varma, D., Wan, X., Civril, F., Cleveland, D.W., Oegema, K., Salmon, E.D., and Desai, A. (2010). Removal of Spindly from microtubule-attached kinetochores controls spindle checkpoint silencing in human cells. Genes Dev 24, 957–971.

Gatlin, J.C., Matov, A., Danuser, G., Mitchison, T.J., and Salmon, E.D. (2010). Directly probing the mechanical properties of the spindle and its matrix. J Cell Biol 188, 481–489.

Grill, S.W., Gonczy, P., Stelzer, E.H., and Hyman, A.A. (2001). Polarity controls forces governing asymmetric spindle positioning in the Caenorhabditis elegans embryo. Nature 409, 630–633.

Guild, J., Ginzberg, M.B., Hueschen, C.L., Mitchison, T.J., and Dumont, S. (2017). Increased lateral microtubule contact at the cell cortex is sufficient to drive mammalian spindle elongation. Mol Biol Cell 28, 1975–1983.

Howell, B.J., McEwen, B.F., Canman, J.C., Hoffman, D.B., Farrar, E.M., Rieder, C.L., and Salmon, E.D. (2001). Cytoplasmic dynein/dynactin drives kinetochore protein transport to the spindle poles and has a role in mitotic spindle checkpoint inactivation. J Cell Biol 155, 1159–1172.

Hueschen, C.L., Galstyan, V., Amouzgar, M., Phillips, R., and Dumont, S. (2019). Microtubule End-Clustering Maintains a Steady-State Spindle Shape. Curr Biol 29, 700–708 e705.

Hueschen, C.L., Kenny, S.J., Xu, K., and Dumont, S. (2017). NuMA recruits dynein activity to microtubule minus-ends at mitosis. eLife 6.

Kapitein, L.C., Peterman, E.J., Kwok, B.H., Kim, J.H., Kapoor, T.M., and Schmidt, C.F. (2005). The bipolar mitotic kinesin Eg5 moves on both microtubules that it crosslinks. Nature 435, 114–118.

Kwon, M., Godinho, S.A., Chandhok, N.S., Ganem, N.J., Azioune, A., Thery, M., and Pellman, D. (2008). Mechanisms to suppress multipolar divisions in cancer cells with extra centrosomes. Genes Dev 22, 2189–2203.

Lancaster, O.M., Le Berre, M., Dimitracopoulos, A., Bonazzi, D., Zlotek-Zlotkiewicz, E., Picone, R., Duke, T., Piel, M., and Baum, B. (2013). Mitotic rounding alters cell geometry to ensure efficient bipolar spindle formation. Dev Cell 25, 270–283.

Le Berre, M., Aubertin, J., and Piel, M. (2012). Fine control of nuclear confinement identifies a threshold deformation leading to lamina rupture and induction of specific genes. Integr Biol (Camb) 4, 1406–1414.

Lecland, N., and Luders, J. (2014). The dynamics of microtubule minus ends in the human mitotic spindle. Nat Cell Biol 16, 770–778.

Ma, N., Tulu, U.S., Ferenz, N.P., Fagerstrom, C., Wilde, A., and Wadsworth, P. (2010). Poleward transport of TPX2 in the mammalian mitotic spindle requires dynein, Eg5, and microtubule flux. Mol Biol Cell 21, 979–988.

Matos, I., Pereira, A.J., Lince-Faria, M., Cameron, L.A., Salmon, E.D., and Maiato, H. (2009). Synchronizing chromosome segregation by flux-dependent force equalization at kinetochores. J Cell Biol 186, 11–26.

Mayer, T.U., Kapoor, T.M., Haggarty, S.J., King, R.W., Schreiber, S.L., and Mitchison, T.J. (1999). Small molecule inhibitor of mitotic spindle bipolarity identified in a phenotype-based screen. Science 286, 971–974.

McIntosh, J.R., Molodtsov, M.I., and Ataullakhanov, F.I. (2012). Biophysics of mitosis. Q Rev Biophys 45, 147–207.

McKinley, K.L., and Cheeseman, I.M. (2017). Large-Scale Analysis of CRISPR/Cas9 Cell-Cycle Knockouts Reveals the Diversity of p53-Dependent Responses to Cell-Cycle Defects. Dev Cell 40, 405–420 e402.

Meijering, E., Dzyubachyk, O., and Smal, I. (2012). Methods for cell and particle tracking. Methods Enzymol 504, 183–200.

Mitchison, T.J., Maddox, P., Gaetz, J., Groen, A., Shirasu, M., Desai, A., Salmon, E.D., and Kapoor, T.M. (2005). Roles of polymerization dynamics, opposed motors, and a tensile element in governing the length of Xenopus extract meiotic spindles. Mol Biol Cell 16, 3064–3076.

Mitra, A., Meissner, L., Gandhimathi, R., Renger, R., Ruhnow, F., and Diez, S. (2020). Kinesin-14 motors drive a right-handed helical motion of antiparallel microtubules around each other. Nat Commun 11, 2565.

Mitra, A., Ruhnow, F., Girardo, S., and Diez, S. (2018). Directionally biased sidestepping of Kip3/kinesin-8 is regulated by ATP waiting time and motor-microtubule interaction strength. Proc Natl Acad Sci U S A 115, E7950–E7959.

Miyamoto, D.T., Perlman, Z.E., Burbank, K.S., Groen, A.C., and Mitchison, T.J. (2004). The kinesin Eg5 drives poleward microtubule flux in Xenopus laevis egg extract spindles. J Cell Biol 167, 813–818.

Mountain, V., Simerly, C., Howard, L., Ando, A., Schatten, G., and Compton, D.A. (1999). The kinesin-related protein, HSET, opposes the activity of Eg5 and cross-links microtubules in the mammalian mitotic spindle. J Cell Biol 147, 351–366.

Neumann, B., Walter, T., Heriche, J.K., Bulkescher, J., Erfle, H., Conrad, C., Rogers, P., Poser, I., Held, M., Liebel, U., et al. (2010). Phenotypic profiling of the human genome by time-lapse microscopy reveals cell division genes. Nature 464, 721–727.

Nitzsche, B., Dudek, E., Hajdo, L., Kasprzak, A.A., Vilfan, A., and Diez, S. (2016). Working stroke of the kinesin-14, ncd, comprises two substeps of different direction. Proc Natl Acad Sci U S A 113, E6582–E6589.

Norris, S.R., Jung, S., Singh, P., Strothman, C.E., Erwin, A.L., Ohi, M.D., Zanic, M., and Ohi, R. (2018). Microtubule minus-end aster organization is driven by processive HSET-tubulin clusters. Nat Commun 9, 2659.

Novak, M., Polak, B., Simunic, J., Boban, Z., Kuzmic, B., Thomae, A.W., Tolic, I.M., and Pavin, N. (2018). The mitotic spindle is chiral due to torques within microtubule bundles. Nat Commun 9, 3571.

Oriola, D., Julicher, F., and Brugues, J. (2020). Active forces shape the metaphase spindle through a mechanical instability. Proc Natl Acad Sci U S A 117, 16154–16159.

Raaijmakers, J.A., and Medema, R.H. (2014). Function and regulation of dynein in mitotic chromosome segregation. Chromosoma 123, 407–422.

Rincon, S.A., Lamson, A., Blackwell, R., Syrovatkina, V., Fraisier, V., Paoletti, A., Betterton, M.D., and Tran, P.T. (2017). Kinesin-5-independent mitotic spindle assembly requires the antiparallel microtubule crosslinker Ase1 in fission yeast. Nat Commun 8, 15286.

Rodenfels, J., Neugebauer, K.M., and Howard, J. (2019). Heat Oscillations Driven by the Embryonic Cell Cycle Reveal the Energetic Costs of Signaling. Dev Cell 48, 646–658 e646.

Roostalu, J., Rickman, J., Thomas, C., Nedelec, F., and Surrey, T. (2018). Determinants of Polar versus Nematic Organization in Networks of Dynamic Microtubules and Mitotic Motors. Cell 175, 796–808 e714.

Saunders, W.S., and Hoyt, M.A. (1992). Kinesin-related proteins required for structural integrity of the mitotic spindle. Cell 70, 451–458.

Sharp, D.J., Yu, K.R., Sisson, J.C., Sullivan, W., and Scholey, J.M. (1999). Antagonistic microtubule-sliding motors position mitotic centrosomes in Drosophila early embryos. Nat Cell Biol 1, 51–54.

Shimamoto, Y., Forth, S., and Kapoor, T.M. (2015). Measuring Pushing and Braking Forces Generated by Ensembles of Kinesin-5 Crosslinking Two Microtubules. Dev Cell 34, 669–681.

Shimamoto, Y., Maeda, Y.T., Ishiwata, S., Libchaber, A.J., and Kapoor, T.M. (2011). Insights into the micromechanical properties of the metaphase spindle. Cell 145, 1062–1074.

Steblyanko, Y., Rajendraprasad, G., Osswald, M., Eibes, S., Jacome, A., Geley, S., Pereira, A.J., Maiato, H., and Barisic, M. (2020). Microtubule poleward flux in human cells is driven by the coordinated action of four kinesins. EMBO J 10.15252/embj.2020105432, e105432.

Sturgill, E.G., and Ohi, R. (2013). Kinesin-12 differentially affects spindle assembly depending on its microtubule substrate. Curr Biol 23, 1280–1290.

Suresh, P., Long, A.F., and Dumont, S. (2020). Microneedle manipulation of the mammalian spindle reveals specialized, short-lived reinforcement near chromosomes. eLife 9.

Takagi, J., Sakamoto, R., Shiratsuchi, G., Maeda, Y.T., and Shimamoto, Y. (2019). Mechanically Distinct Microtubule Arrays Determine the Length and Force Response of the Meiotic Spindle. Dev Cell 49, 267–278 e265.

Tanenbaum, M.E., Macurek, L., Galjart, N., and Medema, R.H. (2008). Dynein, Lis1 and CLIP-170 counteract Eg5-dependent centrosome separation during bipolar spindle assembly. EMBO J 27, 3235–3245.

Tanenbaum, M.E., Macurek, L., Janssen, A., Geers, E.F., Alvarez-Fernandez, M., and Medema, R.H. (2009). Kif15 cooperates with Eg5 to promote bipolar spindle assembly. Curr Biol 19, 1703–1711.

Thévenaz, P., Ruttimann, U.E., and Unser, M. (1998). A pyramid approach to subpixel registration based on intensity. IEEE Trans Image Process 7, 27–41.

Trupinic, M., Ponjavic, I., Kokanovic, B., Barisic, I., Segvic, S., Ivec, A., and Tolic, I.M. (2020). Twist of the mitotic spindle culminates at anaphase onset and depends on microtubule-associated proteins along with external forces. bioRxiv https://doi.org/10.1101/2020.12.27.424486.

van Heesbeen, R.G., Tanenbaum, M.E., and Medema, R.H. (2014). Balanced activity of three mitotic motors is required for bipolar spindle assembly and chromosome segregation. Cell Rep 8, 948–956.

Vanneste, D., Takagi, M., Imamoto, N., and Vernos, I. (2009). The role of Hklp2 in the stabilization and maintenance of spindle bipolarity. Curr Biol 19, 1712–1717.

Vukusic, K., Buda, R., Bosilj, A., Milas, A., Pavin, N., and Tolic, I.M. (2017). Microtubule Sliding within the Bridging Fiber Pushes Kinetochore Fibers Apart to Segregate Chromosomes. Dev Cell 43, 11–23 e16.

Vukusic, K., Buda, R., Ponjavic, I., Risteski, P., and Tolic, I.M. (2019). Chromosome segregation is driven by joint microtubule sliding action of kinesins KIF4A and EG5. bioRxiv https://doi.org/10.1101/863381.

Yajima, J., Mizutani, K., and Nishizaka, T. (2008). A torque component present in mitotic kinesin Eg5 revealed by three-dimensional tracking. Nat Struct Mol Biol 15, 1119–1121.

Yu, C.H., Redemann, S., Wu, H.Y., Kiewisz, R., Yoo, T.Y., Conway, W., Farhadifar, R., Muller-Reichert, T., and Needleman, D. (2019). Central-spindle microtubules are strongly coupled to chromosomes during both anaphase A and anaphase B. Mol Biol Cell 30, 2503–2514.

